# Integrating long-range regulatory interactions to predict gene expression using graph convolutional networks

**DOI:** 10.1101/2020.11.23.394478

**Authors:** Jeremy Bigness, Xavier Loinaz, Shalin Patel, Erica Larschan, Ritambhara Singh

## Abstract

Long-range spatial interactions among genomic regions are critical for regulating gene expression, and their disruption has been associated with a host of diseases. However, when modeling the effects of regulatory factors, most deep learning models either neglect long-range interactions or fail to capture the inherent 3D structure of the underlying genomic organization. To address these limitations, we present GC-MERGE, a **G**raph **C**onvolutional **M**odel for **E**pigenetic **R**egulation of **G**ene **E**xpression. Using a graph-based framework, the model incorporates important information about long-range interactions via a natural encoding of spatial interactions into the graph representation. It integrates measurements of both the spatial genomic organization and local regulatory factors, specifically histone modifications, to not only predict the expression of a given gene of interest but also quantify the importance of its regulatory factors. We apply GC-MERGE to datasets for three cell lines - GM12878 (lymphoblastoid), K562 (myelogenous leukemia), and HUVEC (human umbilical vein endothelial) - and demonstrate its state-of-the-art predictive performance. Crucially, we show that our model is interpretable in terms of the observed biological regulatory factors, highlighting both the histone modifications and the interacting genomic regions contributing to a gene’s predicted expression. We provide model explanations for multiple exemplar genes and validate them with evidence from the literature. Our model presents a novel setup for predicting gene expression by integrating multimodal datasets in a graph convolutional framework. More importantly, it enables interpretation of the biological mechanisms driving the model’s predictions. Available at: https://github.com/rsinghlab/GC-MERGE.

## 1 Introduction

Gene regulation determines the fate of every cell, and its disruption leads to diverse diseases ranging from cancer to neurodegeneration [Krijger and de Laat, 2016, Schoenfelder and Fraser, 2019]. Although specialized cell types – from neurons to cardiac cells – exhibit different gene expression patterns, the information encoded by the linear DNA sequence remains virtually the same in all non-reproductive cells of the body. Therefore, the observed differences in cell type must be encoded by elements extrinsic to sequence, commonly referred to as epigenetic factors. Epigenetic factors found in the local neighborhood of a gene typically include histone marks (also known as histone modifications). These marks are naturally occurring chemical additions to histone proteins that control how tightly the DNA strands are wound around the proteins and the recruitment or occlusion of transcription factors. Recently, the focus of attention in genomics has shifted increasingly to the study of long-range epigenetic regulatory interactions that result from the three-dimensional organization of the genome [Rowley and Corces, 2018]. For example, one early study demonstrated that chromosomal rearrangements, some located as far as 125 kilo-basepairs (kbp) away, disrupted the region downstream of the PAX6 transcription unit causing Aniridia (absence of the iris) and related eye anomalies [Kleinjan et al., 2001]. Thus, chromosomal rearrangement can not only directly affect the expression of proximal genes but can also indirectly affect a gene located far away by perturbing its regulatory (e.g., enhancer-promoter) interactions. This observation indicates that while local regulation of genes is informative, studying long-range gene regulation is critical to understanding cell development and disease. However, experimentally testing for all possible combinations of long-range and short-range regulatory factors for ∼ 20, 000 genes is infeasible given the vast size of the search space. Therefore, computational and data-driven approaches are necessary to efficiently search this space and reduce the number of testable hypotheses.

In recent years, deep learning frameworks have been applied to predict gene expression from histone modifications, and their empirical performance has often exceeded the previous machine learning methods [Cheng et al., 2011, Dong et al., 2012, Karlic et al., 2010]. Among their many advantages, deep neural networks perform automatic feature extraction by efficiently exploring feature space and then finding nonlinear transformations of the weighted averages of those features. This formulation is especially relevant to model complex biological systems since they are inherently nonlinear. For instance, Singh et al. [2016] introduced DeepChrome, which used a convolutional neural network (CNN) to aggregate five types of histone mark ChIP-seq signals in a 10, 000 bp region around the transcription start site (TSS) of each gene. Using a similar setup, they next introduced attention layers to their model [Singh et al., 2017], yielding a comparable performance but with the added ability to visualize feature importance within the local neighborhood of a gene. These methods framed the gene expression problem as a binary classification task in which the gene was either active or inactive. Agarwal and Shendure [2020] introduced Xpresso, a CNN framework that operated on the promoter sequences of each gene and 8 other annotated features associated with mRNA decay to predict steady-state mRNA levels. This model focused primarily on the regression task, such that each prediction corresponded to the logarithm of a gene’s expression. While all the studies mentioned previously accounted for combinatorial interactions among features at the local level, they did not incorporate long-range regulatory interactions known to play a critical role in differentiation and disease [Krijger and de Laat, 2016, Schoenfelder and Fraser, 2019].

Modeling these long-range interactions is a challenging task due to two significant reasons. First, it is difficult to confidently pick an input size for the genomic regions as regulatory elements can control gene expression from various distances. Second, inputting a large region will introduce sparsity and noise into the data, making the learning task difficult. A potential solution to this problem is to incorporate information from long-range interaction networks captured from experimental techniques like Hi-ChIP [Mumbach et al., 2016] and Hi-C [Van Berkum et al., 2010]. These techniques use high-throughput sequencing to measure 3D genomic structure, in which each read pair corresponds to an observed 3D contact between two genomic loci. While Hi-C captures the global interactions of all genomic regions, Hi-ChIP focuses only on spatial interactions mediated by a specific protein. Recently, Zeng et al. [2019b] combined a CNN, encoding promoter sequences, with a fully connected network using Hi-ChIP datasets to predict gene expression values. The authors then evaluated the relative contributions of the promoter sequence and promoter-enhancer submodules to the model’s overall performance. While this method incorporated long-range interaction information, its use of HiChIP experiments narrowed this information to spatial interactions facilitated by H3K27ac and YY1. Furthermore, CNN models can only capture local topological patterns instead of modeling the underlying spatial structure of the data, thus limiting interpretation to local sequence features.

To address these issues, we developed a **G**raph **C**onvolutional **M**odel of **E**pigentic **R**egulation of **G**ene **E**xpression (GC-MERGE), a graph-based deep learning framework that integrates 3D genomic data with histone mark signals to predict gene expression. Figure 1 provides a schematic of our overall approach. Unlike previous methods, our model incorporates genome-wide interaction frequencies of the Hi-C data by encoding it via a graph convolutional network (GCN), thereby capturing the underlying spatial structure. GCNs are particularly well-suited to representing spatial relationships, as a Hi-C map can be represented as an adjacency matrix of an undirected graph *G* ∈ {*V, E*}. Here, *V* nodes represent the genomic regions and *E* edges represent their interactions. Our formulation leverages information from both local as well as distal regulatory factors that control gene expression. While some methods use a variety of other features, such as promoter sequences or ATAC-seq levels [Agarwal and Shendure, 2020, Dong et al., 2012, Zeng et al., 2019b], we focus our efforts solely on histone modifications and extract their relationship to the genes. We show that our model provides state-of-the-art performance for the gene expression prediction tasks even with this simplified set of features for three difference cell lines - GM12878 (lymphoblastoid), K562 (myelogenous leukemia), and HUVEC (human umbilical vein endothelial).

**Figure 1:**
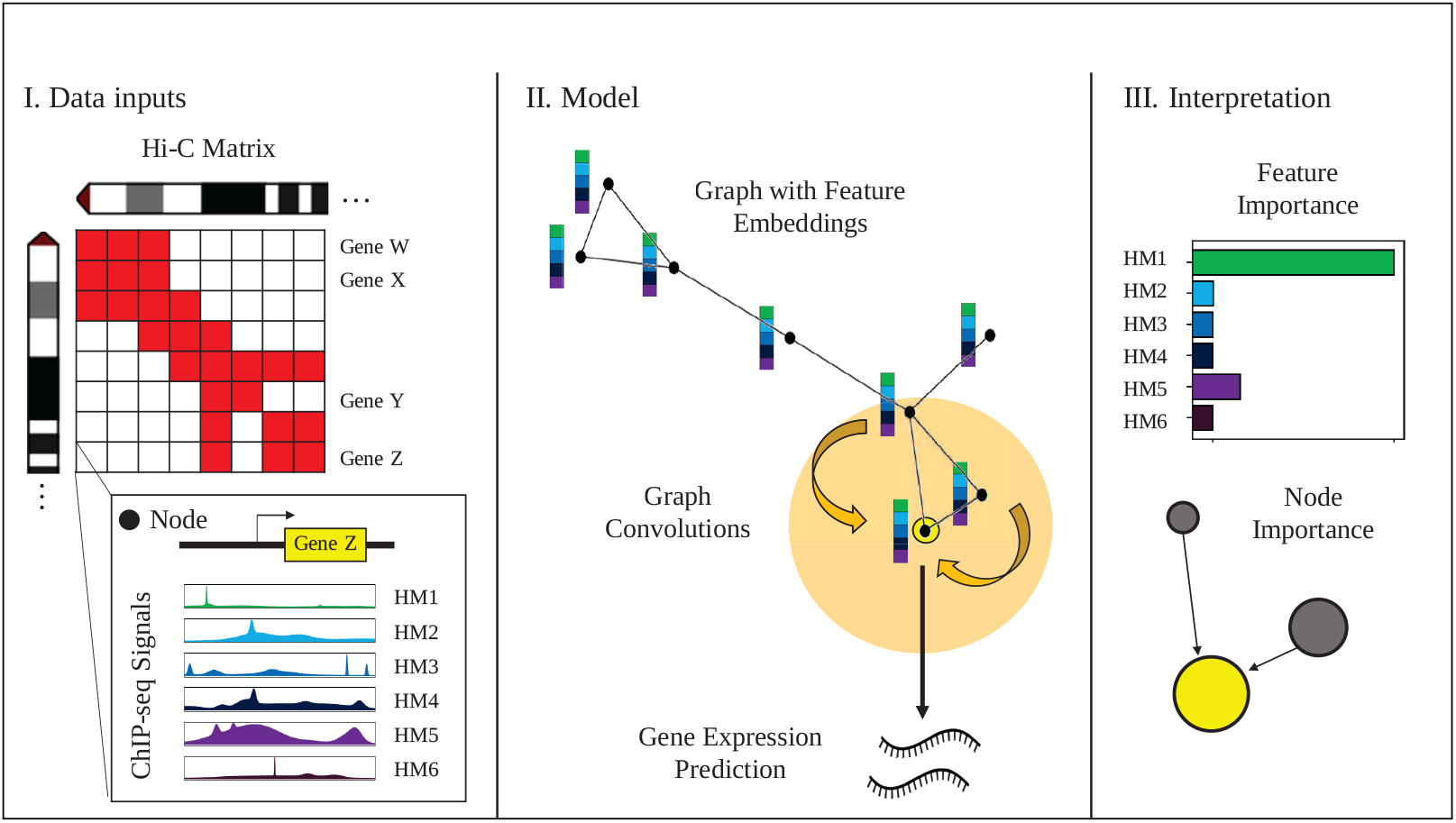
Overview of GC-MERGE. Our framework integrates local histone mark (HM) signals and long-range spatial interactions to predict and understand gene expression. (I) Inputs to the model include Hi-C maps for each chromosome, with the binned chromosomal regions corresponding to nodes in the graph, and the average ChIP-seq readings of six core histone marks in each region, which constitute the initial feature embedding of the nodes. (II) For nodes corresponding to regions containing a gene, the model performs repeated graph convolutions over the neighboring nodes to yield either a binarized class prediction of gene expression activity (either active or inactive) or a continuous, real-valued prediction of expression level. (III) Finally, explanations for the model’s predictions for any gene-associated node can be obtained by calculating the importance scores for each of the features and the relative contributions of neighboring nodes. Therefore, the model provides biological insight into the pattern of histone marks and the genomic interactions that work together to predict gene expression.

A significant contribution of our work is to enable researchers to determine which regulatory interactions – local or distal – contribute towards the gene’s expression prediction and which histone marks are involved in these interactions. This information can suggest promising hypotheses and guide new research directions by making the model’s predictive drivers more transparent. To that effect, we adapt a recent model explanation approach specifically for GCNs known as GNNExplainer [Ying et al., 2019], which quantifies the relative importance of the nodes and edges in a graph that drive the output prediction. We integrate this method within our modeling framework to highlight the important histone modifications (node features) and the important long-range interactions (edges) that contribute to a particular gene’s predicted expression. To validate the model’s explanations, we use two high-throughput experimental studies [Jung et al., 2019, Fulco et al., 2019] that identify significant regulatory interactions. While existing methods [Singh et al., 2016, 2017, Agarwal and Shendure, 2020, Zeng et al., 2019b] can provide feature-level interpretations (important histone modifications or sequences), the unique modeling of Hi-C data as a graph allows GC-MERGE to provide additional edge-level interpretations (important local and global interactions in the genome). Table 1 places the proposed framework among state-of-the-art deep learning models and lists each model’s properties.

**Table 1:** Comparison of the properties of previous deep learning models predicting gene expression with GC-MERGE. The proposed method integrates local and long-range regulatory interactions, capturing the underlying 3D genomic spatial structure as well as highlighting both the critical node-level (histone modifications) and edge-level (genomic interactions) features.

| Computational Study | Inputs |  |  | Interpretation |  |
| --- | --- | --- | --- | --- | --- |
|  | Local Histone Marks | Additional Features<br>(e.g. promoter sequence) | Long-range Interactions | Feature-level Interpretation | Edge-level Interpretation |
| DeepChrome | X |  |  | X |  |
| AttentiveChrome | X |  |  | X |  |
| Xpresso |  | X |  | X |  |
| DeepExpression |  | X | X | X |  |
| GC-MERGE | X |  | X | X | X |

## 2 Methods

### 2.1 Graph convolutional networks (GCNs)

Graph convolutional networks (GCNs) are a generalization of convolutional neural networks (CNN s) to graph-based relational data that is not natively structured in Euclidean space [Liu and Zhou, 2020]. Due to the expressive power of graphs, GCNs have been applied across a wide variety of domains, including recommender systems [Jin et al., 2020] and social networks [Qiu et al., 2018]. The prevalence of graphs in biology has made these models a popular choice for tasks like characterizing protein-protein interactions [Yang et al., 2020], predicting chromatin signature profiles [Lanchantin and Qi, 2020], and inferring the chemical reactivity of molecules for drug discovery [Sun et al., 2020].

We use the GraphSAGE formulation [Hamilton et al., 2017] as our GCN for its relative simplicity and its capacity to learn generalizable, inductive representations not limited to a specific graph. The input to the model is represented as a graph *G* ∈ {*V, E*}, with nodes *V* and edges *E*, and a corresponding adjacency matrix A ∈ ℝ*^N^* ^×^*^N^* [Liu and Zhou, 2020], where *N* is the number of nodes. For each node *v*, there is also an associated feature vector *x_v_*. The goal of the network is to learn a state embedding 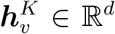 for *v*, which is obtained by aggregating information over *v*’s neighborhood *K* times, where *d* is the dimension of the embedding vector. This new state embedding is then fed through a fully-connected network to produce an output 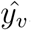, which can then be applied to downstream classification or regression tasks.

Within this modeling framework, the first step is to initialize each node with its input features. In our case, the feature vector *x_v_* ∈ ℝ*^m^* is obtained from the ChIP-seq signals corresponding to the six (*m* = 6) core histone marks (H3K4me1, H3K4me3, H3K9me3, H3K36me3, H3K27me3, and H3K27ac) in our dataset:

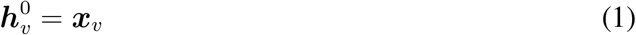

Next, to transition from the (*k* − 1)*^th^* layer to the *k^th^* hidden layer in the network for node *v*, we apply an aggregation function to the neighborhood of each node. This aggregation function is analogous to a convolution operation over regularly structured Euclidean data such as images. While standard convolution function operates over a grid and represents a pixel as a weighted aggregation of its neighboring pixels, in an analogous manner, a graph convolution performs this operation over the neighbors of a node in a graph. In our case, the aggregation function calculates the mean of the neighboring node features:

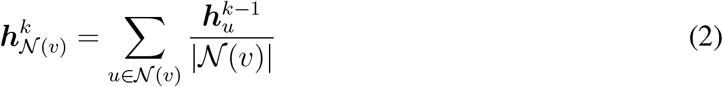

Here, *N* (*v*) represents the adjacency set of node *v*. We update the node’s embedding by concatenating the aggregation with the previous layer’s representation to retain information from the original embedding. Next, just as done in a standard convolution operation, we take the matrix product of this concatenated representation with a learnable weight matrix to complete the weighted aggregation step. Finally, we apply a non-linear activation function, such as ReLU, to capture the higher-order non-linear interactions among the features:

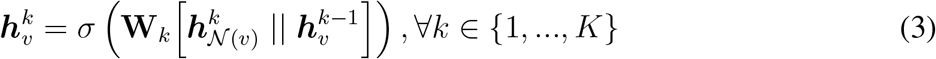

Here, ‖ represents concatenation, *σ* is a non-linear activation function, and *W_k_* is a learnable weight parameter. After this step, each node is assigned a new embedding. After *K* iterations, the node embedding encodes information from the neighbors that are *K*-hops away from that node:

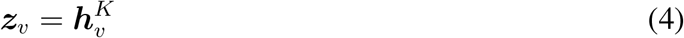

Here, z*_v_* is the final node embedding after *K* iterations.

GC-MERGE is a flexible framework that can formulate gene expression prediction as both a classification and a regression task. For the classification task, we feed the learned embedding z*_v_* into a fully connected network and output a prediction 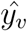 for each target node using a *Softmax* layer to compute probabilities for each class *c* and then take the *argmax*. Here, class *c* ∈ {0, 1} corresponds to whether the gene is either off/inactive (*c* = 0) or on/active (*c* = 1). We use the true binarized gene expression value *y_v_* ∈ {0, 1} by thresholding the expression level relative to the median as the target predictions, consistent with other studies [Singh et al., 2016, 2017]. For the loss function, we minimize the negative log likelihood (NLL) of the log of the *Softmax* probabilities. For the regression task, we feed *z_v_* into a fully connected network and output a prediction 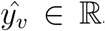, representing a real-valued expression level. We use the mean squared error (MSE) as the loss function. For both tasks, the model architecture is summarized in Figure 2 and described in further detail in Supplemental Section S1.1.

**Figure 2:**
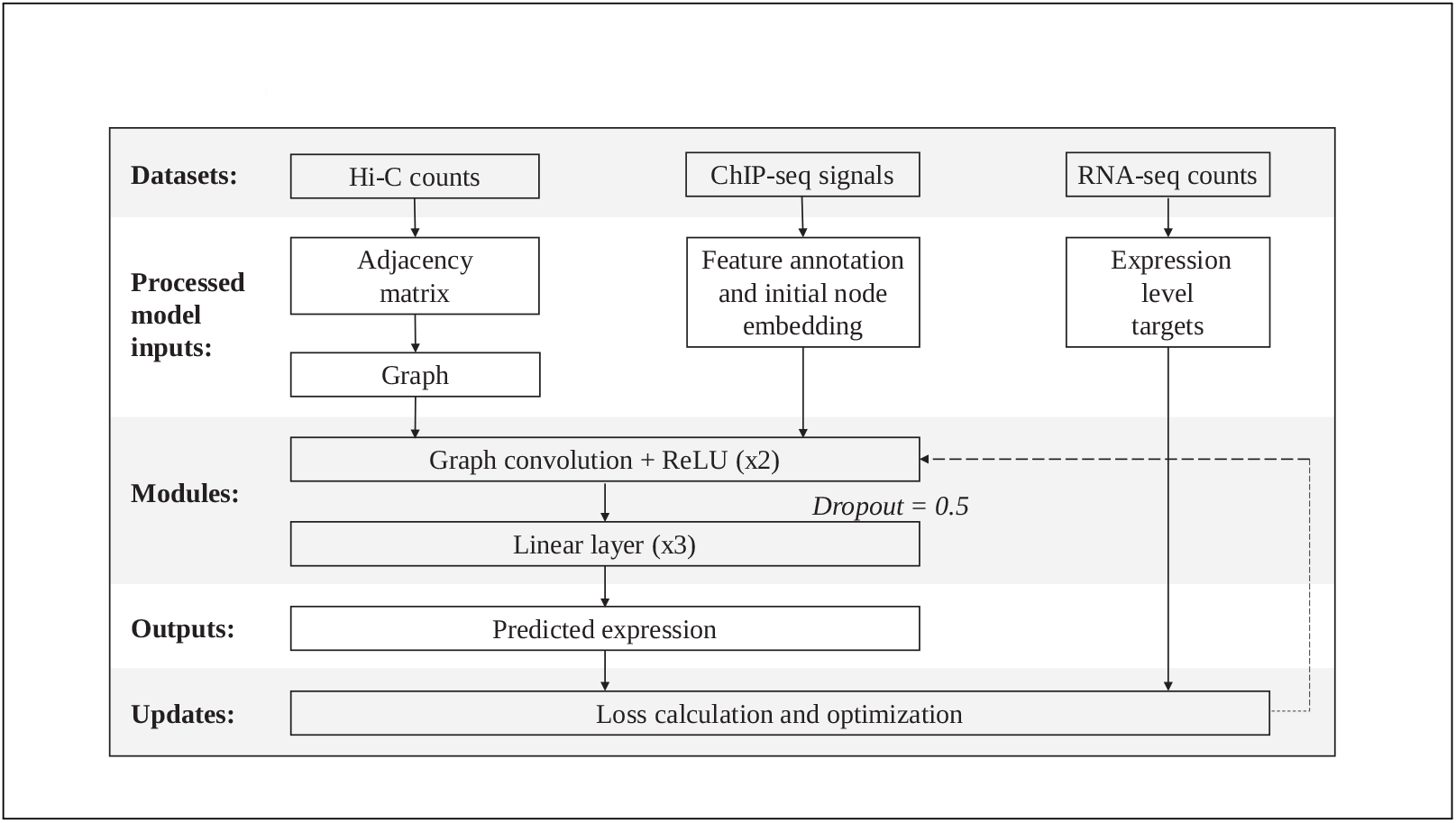
Overview of the GCNN model architecture. The datasets used in our model are Hi-C maps, ChIP-seq signals, and RNA-seq counts. A binarized adjacency matrix (A ∈ ℝ*^N^* ^×^*^N^* ) is produced from the Hi-C maps by subsampling from the Hi-C matrix. The nodes *v* in the graph are annotated with features from the ChIP-seq datasets (x_v_). Two graph convolutions, each followed by ReLU, are performed. The output from here is fed into a dropout layer (probability = 0.5), followed by a linear module comprised of three dense layers, in which ReLU follows the first two layers. For the classification model, the output is fed through a *softmax* layer, and then the *argmax* is taken to make the final prediction (*y_v_* ). For the regression model, the final output represents the base-10 logarithm of the expression level (with a pseudocount of 1).

### 2.2 Interpretation of GC-MERGE

Although a model’s architecture is integral to its performance, just as important is understanding how the model arrives at its predictions. Neural networks, in particular, have sometimes been criticized for being “black box” models, such that no insight is provided into how the model operates. Most graph-based interpretability approaches either approximate models with simpler models whose decisions can be used for explanations [Ribeiro et al., 2016] or use an attention mechanism to identify relevant features in the input that guide a particular prediction [Veličković et al., 2017]. In general, these methods, along with gradient-based approaches [Simonyan et al., 2013, Sundararajan et al., 2017] or DeepLift [Shrikumar et al., 2017], focus on the explanation of important node features and do not incorporate the structural information of the graph. However, a recent method called *Graph Neural Net Explainer* (or GNNExplainer) [Ying et al., 2019], given a trained GCN, can identify a small subgraph as well as a small subset of features that are crucial for a particular prediction.

We adapt the GNNExplainer method and integrate it into our classifier framework. GNNEx-plainer maximizes the mutual information between the probability distribution of the model’s class predictions over all nodes and the probability distribution of the class predictions for a particular node conditioned on some fractional masked subgraph of neighboring nodes and features. Subject to regularization constraints, it jointly optimizes the fractional node and feature masks, determining the extent to which each element informs the prediction for a particular node.

Specifically, given a node *v*, the goal is to learn a subgraph *G_s_* ⊆ *G* and a feature mask *X_s_* = {*x_j_* | *v_j_* ∈ *G_s_*} that contribute the most toward driving the full model’s prediction of 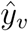 To achieve this objective, the algorithm learns a mask that maximizes the mutual information (MI) between the original model and the masked model. Mathematically, this objective function is as follows:

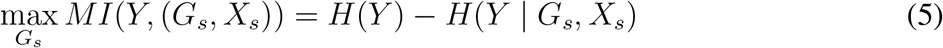

where *H* is the entropy of a distribution. Since this is computationally intractable with an exponential number of graph masks, GNNExplainer optimizes the following quantity using gradient descent:

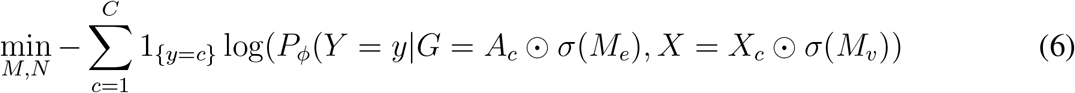

where *c* represents the class, *A_c_* represents the adjacency matrix of the computation graph, *M_e_* represents the subgraph mask on the edges, and *M_v_* represents the node feature mask. The importance scores of the nodes and features are obtained by applying the sigmoid function to the subgraph edges and node feature masks, respectively. Finally, the element-wise entropies of the masks are calculated and added as regularization terms into the loss function. Therefore, in the context of our model, GNNExplainer learns which genomic interactions (via the subgraph edge mask) and which histone modifications (via the node feature mask) are most critical to driving the model’s predictions.

## 3 Experimental Setup

### 3.1 Overview of the datasets

GC-MERGE requires the following information: (1) Interactions between the genomic regions (Hi-C contact maps); (2) Histone mark signals representing the regulatory signals (ChIP-seq measurements); (3) Expression levels for each gene (RNA-seq measurements). Thus, for each gene in a particular region, the first two datasets are the inputs into our proposed model, whereas gene expression is the predicted target.

Being consistent with previous studies [Singh et al., 2016, 2017], we first formulate the prediction problem as a classification task. However, as researchers may be interested in predicting exact expression levels, we also extend the predictive capabilities of our model to the regression setting. For the classification task, we binarize the gene expression values as either 0 (low expression) or 1 (high expression) using the median as the threshold, as done in previous studies [Cheng et al., 2011, Singh et al., 2016, 2017, Zeng et al., 2019b]. For the regression task, we take the base-10 logarithm of the gene expression values with a pseudo-count of 1.

We focused our experiments on three human cell lines from Rao et al. [2014]: (1) GM12878, a lymphoblastoid cell line with a normal karyotype, (2) K562, a myelogenous leukemia cell line, and (3) HUVEC, a human umbilical vein endothelial cell line. For each of these cell lines, we accessed RNA-seq expression and ChIP-Seq signal datasets for six uniformly profiled histone marks from the REMC repository [Roadmap Epigenomics Consortium, 2015]. These histone marks include (1) H3K4me1, associated with enhancer regions; (2) H3K4me3, associated with promoter regions; (3) H3K9me3, associated with heterochromatin; (4) H3K36me3, associated with actively transcribed regions; (5) H3K27me3, associated with polycomb repression; and (6) H3K27ac, also associated with enhancer regions. We chose these marks because of the wide availability of the relevant data sets as well as for ease of comparison with previous studies [Singh et al., 2016, 2017, Zeng et al., 2019b]. In addition, these six core histone marks are the same set of features used in the widely-cited 18-state ChromHMM model [Ernst and Kellis, 2017], which associates histone mark signatures with chromatin states.

### 3.2 Graph construction and data integration

Our main innovation is formulating the graph-based prediction task to integrate two very different data modalities (histone mark signals and Hi-C interaction frequencies). We represented each genomic region with a node (*v*) and connected an edge (*e*) between it and the nodes corresponding to its neighbors (bins with non-zero entries in the adjacency matrix) to construct the graph (*G* ∈ {*V, E*}, with nodes *V* and edges *E*). For chromosome capture data, we used previously published Hi-C maps at 10 kilobase-pair (kbp) resolution for all 22 autosomal chromosomes [Rao et al., 2014]. We obtained an *N* × *N* symmetric matrix, where each row or column represents a 10 kb chromosomal region. Therefore, each bin count corresponds to the interaction frequency between the two respective genomic regions. Next, we applied VC-normalization on the Hi-C maps. In addition, because chromosomal regions located closer together will contact each other more frequently than regions located farther away simply due to chance (rather than due to biologically significant effects), we made an additional adjustment for this background effect. Following Sobhy et al. [2019], we determined the distance between the regions corresponding to each row and column. Then, for all pairs of interacting regions located the same distance away, we calculated the median of the bin counts along each diagonal of the *N* x*N* matrix and used this as a proxy for the background. Finally, for each bin, we subtracted the appropriate median and discarded any negative values. We converted all non-zero values to 1, thus obtaining the binary adjacency matrix for our model (A ∈ ℝ*^N^* ^×^*^N^* ).

Due to the large size of the Hi-C graph, we subsampled neighbors to form a subgraph for each node we fed into the model. While there are methods to perform subsampling on large graphs using a random node selection approach (e.g., Zeng et al. [2019a]), we used a simple strategy of selecting the top *j* neighbors with the highest Hi-C interaction frequency values. We empirically selected the value of *j* = 10 for the number of neighbors. Increasing the size of the subsampled neighbor set (i.e., *j* = 20) did not improve the performance further, as shown in Supplementary Figure S1.

To integrate the Hi-C datasets with the RNA-seq and ChIP-seq datasets, we obtained the average ChIP-seq signal for each of the six core histone marks over the 10 kbp chromosomal region corresponding to each node. In this way, we associated a feature vector of length six with each node (x_v_ ∈ ℝ^6^). For assigning an output value to the node, we took each gene’s transcriptional start site (TSS) and assigned its expression value to the node corresponding to the chromosomal region with its TSS as output (*y_v_* ). If multiple genes were assigned to the same node, we took the median of the expression levels, i.e., the median of all the values corresponding to the same node. Given our framework, we could allot the output gene expression to only a subset of nodes that contained gene TSSs while aiming to use histone modification signals from all the nodes. Therefore, to enable training with such a unique setting, we applied a mask during the training phase so that the model made predictions only on nodes with assigned gene expression values. Note that the graph convolution operation used information from all the related nodes but made predictions on the subset of nodes with output values.

The overall size of our data set consisted of 279, 606 total nodes and 16, 699 gene-associated nodes for GM12878, 279, 601 total nodes and 16, 690 gene-associated nodes for K562, and 279, 59-8 total nodes and 16, 681 gene-associated nodes for HUVEC. When running the model on each cell line, we assigned 70% of the gene-associated nodes to the training set, 15% to the validation set, and 15% to the testing set. Then, we performed hyperparameter tuning using the training and validation sets and report performance on the independent test set. The details of the hyperparameter tuning are provided in Supplementary section S1.2.

### 3.3 Baseline models

We compared GC-MERGE with the following deep learning baselines for gene expression prediction both the classification and regression tasks:

• **Multi-layer perceptron (MLP)**: A neural network comprised of three fully connected layers.

• **Shuffled neighbor model**: GC-MERGE applied to shuffled Hi-C matrices, such that the neighbors of each node are randomized. We include this baseline to see how the performance of GCN is affected when the provided spatial information is random.

• **Convolutional neural network (CNN)**: A convolutional neural network based on DeepChrome [Singh et al., 2016]. This model takes 10 kb regions corresponding to the genomic regions demarcated in the Hi-C data and subdivides each region into 100 bins. Each bin is associated with six channels, corresponding to the ChIP-seq signals of the six core histone marks used in the present study. A standard convolution is applied to the channels, followed by a fully connected network.

For the regression task, the range of the outputs is the set of continuous real numbers. For the classification task, a *Softmax* function is applied to the model’s output to yield a binary prediction. None of the baseline methods incorporate spatial information. Therefore, they only process histone modification information from the regions whose gene expression is being predicted. In contrast, GC-MERGE solves a more challenging task by processing information from the neighboring regions as well.

For the CNN baseline, genomic regions are subdivided into smaller 100-bp bins, consistent with Singh et al. [2016]. However, GC-MERGE and the baselines other than the CNN average the histone modification signals over the entire 10 kb region. We also implemented GC-MERGE on higher resolution ChIP-seq datasets (1000-bp bins), which we fed through a linear embedding module to form features for the Hi-C nodes. We did not observe an improvement in the performance for the high-resolution input (Supplemental Figure S2).

Additionally, we compared our results to the published results of two other recent deep learning methods, Xpresso by Agarwal and Shendure [2020] and DeepExpression by Zeng et al. [2019b], when such comparisons were possible, although in some cases the experimental data sets were unavailable or the code provided did not run.

### 3.4 Evaluation metrics

For the classification task, we evaluated model performance by using two metrics: the area under the receiver operating characteristic curve (AUROC) and the area under the precision-recall curve (AUPR). For the regression task, we calculated the Pearson correlation coefficient (PCC), which quantifies the correlation between the true and predicted gene expression values in the test set.

## 4 Results

### 4.1 GC-MERGE gives state-of-the-art performance for the gene expression prediction task

We evaluate GC-MERGE and the baseline models on both the classification and regression tasks for the GM12878, K562, and HUVEC cell lines. As earlier studies formulated the problem as a classification task [Singh et al., 2016, 2017, Zeng et al., 2019b], we first apply GC-MERGE to make a binary prediction of whether each gene is active or inactive. In Figure 3(a), we show that our model’s performance is an improvement over all other alternatives, achieving 0.91, 0.92, and 0.90 AUROC scores. We also measure model performance using the AUPR score and achieve similar results (Supplementary Figure S3). For the K562 cell line, we note that the performance of GC-MERGE (AUROC = 0.92) is similar to that reported for DeepExpression (AUROC = 0.91) by Zeng et al. [2019b], a CNN model that uses promoter sequence data as well as spatial information from H3K27ac and YY1 Hi-ChIP experiments. We could not compare to DeepExpression for the GM12878 and HUVEC cell lines as the experimental data sets were unavailable. For the Xpresso framework presented in Agarwal and Shendure [2020], a CNN model that uses promoter sequence and 8 features associated with mRNA decay to predict gene expression, the task is formulated as a regression problem, so no comparisons could be made for the classification setting.

**Figure 3:**
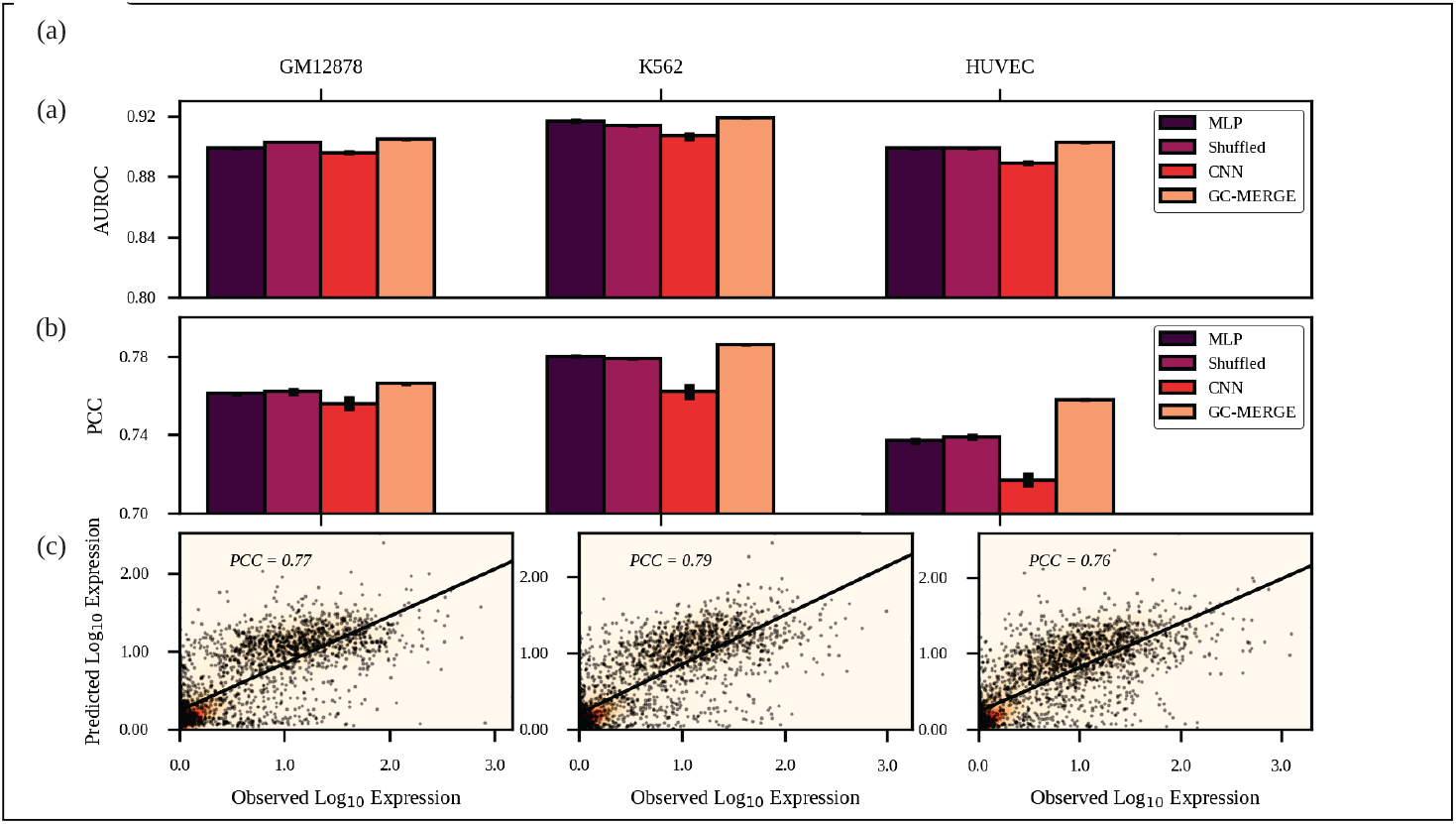
Comparison of AUROC and PCC scores for all models. GC-MERGE gives state-of-the-art performance for both the classification and the regression task. For each reported metric, we take the average of ten runs and denote the standard deviation by the error bars on the graph. (a) For the classification task, the AUROC metrics for GM12878, K562, and HUVEC were 0.91, 0.92, and 0.90, respectively. For each of these cell lines, GC-MERGE improves prediction performance over other baselines. (b) For the regression task, GC-MERGE obtains PCC scores of 0.77, 0.79, and 0.76 for GM12878, K562, and HUVEC, respectively. These scores are better than the respective baselines. (c) Scatter plots of the logarithm of the predicted expression values versus the true expression values are shown for all three cell lines.

With respect to the regression task, Figure 3(b) compares our model’s performance with the baselines and Figure 3(c) shows the predicted versus true gene expression values for GC-MERGE. For GM12878, the Pearson correlation coefficient of GC-MERGE predictions (PCC = 0.77) is better than the other baselines. Furthermore, we note that our model performance also compares favorably to numbers reported for Xpresso (PCC ≈ 0.65) [Agarwal and Shendure, 2020]. For K562, GC-MERGE again outperforms all alternative baseline models (PCC = 0.79). In addition, GC-MERGE performance also exceeds that of Xpresso (PCC ≈ 0.71) [Agarwal and Shendure, 2020] as well as DeepExpression (PCC = 0.65) [Zeng et al., 2019b]. Our model gives better performance (PCC = 0.76) relative to the baselines for HUVEC as well. Neither Xpresso nor DeepExpression studied this cell line. While the metrics presented for GC-MERGE are not directly comparable to the reported numbers for Xpresso and DeepExpression, it is encouraging to see that they are in the range of these state-of-the-art results. An interesting observation here is that the shuffled baseline behaves very similar to the MLP. We hypothesize that the GCN models will most likely ignore the random interaction information and focus on the histone modification signals to make the predictions.

Furthermore, compared to the CNN and MLP baselines, our results suggest that including spatial information can improve gene expression predictive performance over methods that solely use local histone mark features as inputs. We want to emphasize that while all models can predict reasonably well, only GC-MERGE can model the spatial information across multiple genomic regions (including those not associated with the gene) with histone modifications to predict gene expression. Therefore, a state-of-the-art performance on this challenging task indicates that the model can leverage multimodal data sets to learn the relevant connections. An important aim is to go beyond the prediction task and extract these learned relationships from the model. Thus, we present GC-MERGE as a hypothesis driving tool for understanding epigenetic regulation.

### 4.2 Interpretation of GC-MERGE highlights relevant long-range interactions and histone modification profiles

To determine the underlying biological factors driving the model’s predictions, we integrate the GNNExplainer method [Ying et al., 2019], designed for classification tasks, into our modeling framework. Once trained, we show that GC-MERGE can determine which spatial interactions and histone marks are most critical to a gene’s expression prediction. We validate our approach using two experimental data sets that identify interactions of regulatory elements. The first data set is drawn from an analytical study by Jung et al. [2019], which uses promoter capture Hi-C to identify candidate regulatory elements that interact with promoters of interest in conjunction with eQTL expression levels and other epigenetic signals. The second functional characterization study by Fulco et al. [2019], introduces a new experimental technique called CRISPRi-FlowFISH, in which candidate regulatory elements are perturbed, and the effects on the expression of specific genes of interest are measured.

For the promoter capture Hi-C data [Jung et al., 2019], we examined GM12878, a lymphoblastoid cell line, and selected four exemplar genes that are among the most highly expressed in our data set: SIDT1, AKR1B1, LAPTM5, and TOP2B. Brief descriptions of the genes are included in Supplemental Section S2 and the chromosomal coordinates and corresponding node identifiers for each gene can be found in Supplemental Table S2. In Figure 4(a), we show that for SIDT1, the nodes that are ranked as the top three by importance score (indicated by the size of the node) correspond to known regulatory regions. In addition, we plot the importance scores assigned to the histone marks (node features) that are most consequential in determining the model’s predictions. The bar graph shows that H3K4me3 is the most important feature in determining the model’s prediction. This histone mark profile has been associated with regions flanking transcription start sites (TSS) in highly expressed genes [Ernst and Kellis, 2017]. We report similar results for AKR1B1 (Figure 4(b)), where the node ranked as the most important corresponds to a confirmed regulatory region and TOP2B (Figure 4(d)), where two of the most important nodes correspond to regulatory regions. For LAPTM5, shown in Figure 4(c), the top-ranked node corresponds to a validated regulatory region. For the histone importance score profile, the feature deemed most important is H3K27ac. This histone mark has been associated with the promoter regions of highly expressed genes as well as active enhancer regions [Ernst and Kellis, 2017].

**Figure 4:**
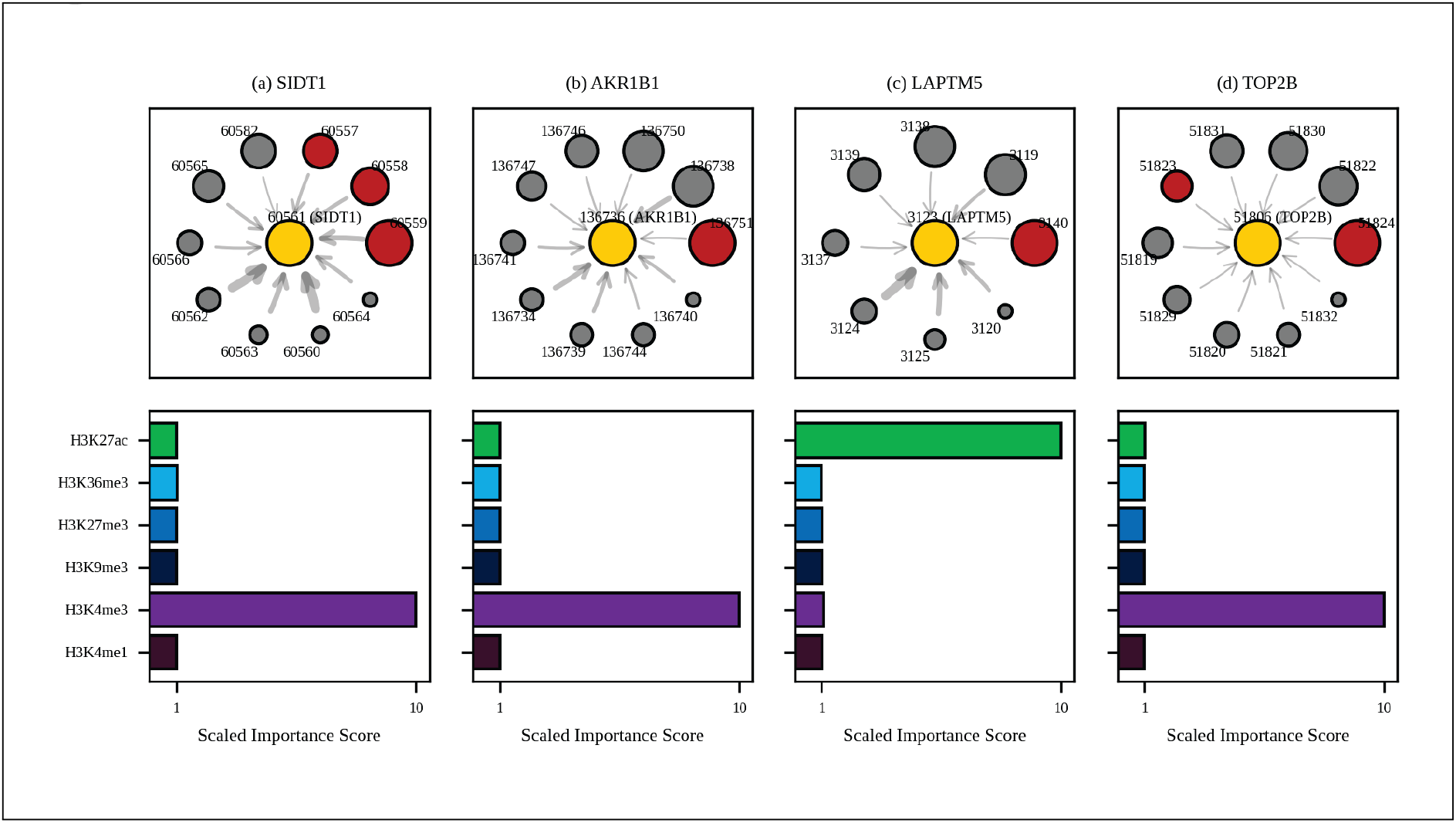
Model explanations for exemplar genes validated by promoter-capture Hi-C. Top: For (a) SIDT1, designated as node 60561 (yellow circle), the subgraph of neighbor nodes is displayed. The size of each neighbor node correlates with its predictive importance as determined by GNNExplainer. Nodes in red denote regions corresponding to known enhancer regions regulating SIDT1 [Jung et al., 2019] (note that multiple interacting fragments can be assigned to each node, see Supplemental Table S3). All other nodes are displayed in gray. The thickness of each edge is inversely correlated with the genomic distance between each neighbor node and the central node, such that thicker edges indicate neighbor nodes that are closer in sequence-space to the gene of interest. Nodes with importance scores corresponding to outliers have been removed for clarity. **Bottom**: The scaled feature importance scores for each of the six core histone marks used in this study are shown in the bar graph. Results also presented for (b) AKR1B1, (c) LAPTM5, and (d) TOP2B.

For the CRISPRi-FlowFISH data set [Fulco et al., 2019], we again highlight four exemplar genes: BAX, HNRNPA1, PRDX2, and RAD23A. Descriptions of each of these genes can be found in Supplementary Section S2 and the gene coordinates and corresponding node IDs can be found in Supplementary Table S2. For BAX, shown in Figure 5(a), the two top-ranked nodes by importance score correspond to functional enhancer regions. The histone mark importance scores pinpoint the H3K4me3 mark as most critical to the model’s predictions. For HNRNPA1 (Figure 5(b)), two out of the three highest-ranked nodes correspond to regulatory regions. The histone marks most important to the model’s predictions are H3K36me3, H3K27ac, and H3K4me3. This chromatin signature is indicative of genic enhancer regions [Ernst and Kellis, 2017]. For PRXD2 (Figure (5(c)), the top two nodes by importance correspond to functional enhancer regions, and the histone mark importance scores indicate that H3K27ac and H3K4me3 play crucial roles in driving the gene’s predicted expression. For RAD23A (Figure 5(d)), the top two nodes again correspond to experimentally validated regulatory regions. From the histone mark importance profile, it can be seen that H3K27ac plays an influential role.

**Figure 5:**
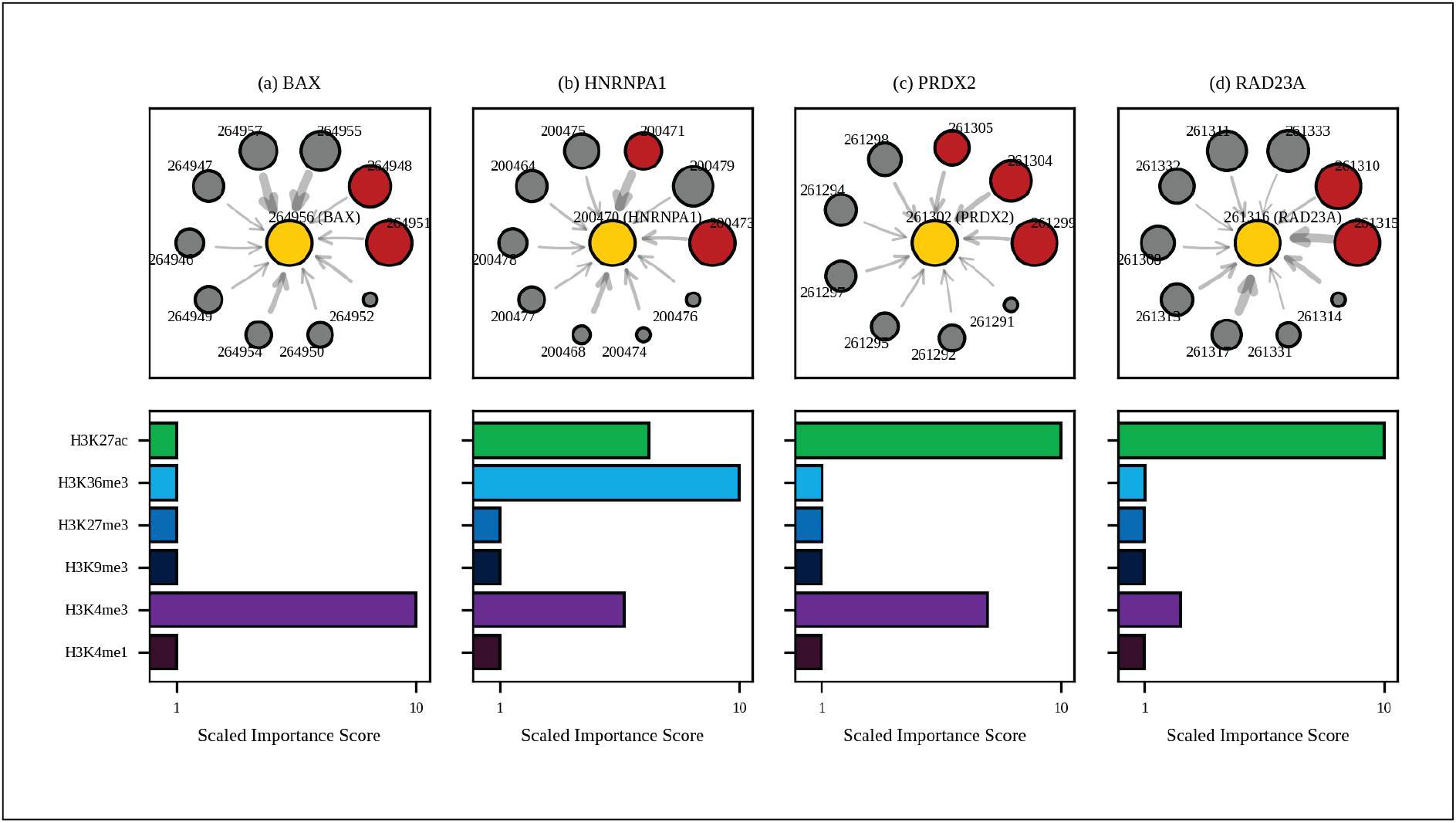
Model explanations for exemplar genes validated by CRISPRi-FlowFISH. Top: For (a) BAX, designated as node 264956 (yellow circle), the subgraph of neighbor nodes is displayed. The size of each neighbor node correlates with its predictive importance as determined by GNNExplainer. Nodes in red denote regions corresponding to known enhancer regions regulating BAX [Fulco et al., 2019] (note that multiple interacting fragments can be assigned to each node, see Supplemental Table S3). All other nodes are displayed in gray. The thickness of each edge is inversely correlated with the genomic distance between each neighbor node and the central node, such that thicker edges indicate neighbor nodes that are closer in sequence-space to the gene of interest. Nodes with importance scores corresponding to outliers have been removed for clarity. Bottom: The scaled feature importance scores for each of the six core histone marks used in this study are shown in the bar graph. Results also presented for (b) HNRNPA1, (c) PRDX2, and (d) RAD23A.

Both H3K4me3 and H3K27ac are active *cis*-regulatory elements used to deduce the enhancer-promoter interactions [Salviato et al., 2021], and, interestingly, interpretation of GC-MERGE highlights these histone marks out of the six chosen for this study.

To confirm that the node importance scores obtained from GNNExplainer do not merely reflect the relative magnitudes of the Hi-C counts or the distances between genomic regions, we investigated the relationships among the Hi-C counts, genomic distances, and scaled importance scores. We observe that the scaled importance scores do not correlate to the Hi-C counts or the pairwise genomic distances. For instance, for SIDT1 (Supplemental Figure S4(a) and Supplemental Table S4), the three experimentally validated interacting nodes have importance scores ranking among the highest (10.0, 6.6 and 5.7). However, they do not correspond to the nodes with the most Hi-C counts (413, 171, and 155 for each of the three known regulatory regions, while the highest count is 603). In addition, these nodes are located 20, 30, and 40 kbp away from the gene region – distances which are characteristic of distal enhancers [Dekker and Misteli, 2015] – while other nodes at the same or closer distances do not have promoter-enhancer interactions. For LAPTM5 (Supplemental Figure S4(c) and Supplemental Table S4), the node with the highest importance score has an experimentally confirmed interaction and is located 170 kbp away from the gene region. We perform similar analysis for all of the other exemplar genes (Supplemental Figure S4 and Supplemental Table S4). Therefore, we show that by modeling the histone modifications and the spatial configuration of the genome, GC-MERGE infers connections that can serve as important hypothesis-driving observations for gene regulatory experiments.

## 5 Discussion

We present GC-MERGE, a graph-based deep learning model, which integrates both local and long-range epigenetic data in a GCN framework to predict gene expression and explain its chief drivers. We demonstrate the model’s state-of-the-art performance for the gene expression prediction task, outperforming the baselines on the GM12878, K562, and HUVEC cell lines. We also determine the relative contributions of histone modifications and genomic interactions for multiple exemplar genes, showing that our model recapitulates known experimental results in a biologically interpretable manner.

For future work, we anticipate applying our model to additional cell lines as high-quality Hi-C data sets become available. Another avenue of particular importance would be to develop more robust methods for interpreting GCNs. For example, while the GNNExplainer model is a theoretically sound framework and yields an unbiased estimator for the importance scores of the subgraph nodes and features, there is variation in the interpretation scores generated over multiple runs. Furthermore, with larger GCNs, the optimization function utilized in GNNExplainer is challenging to minimize in practice. The importance scores converge with little differentiation for some iterations, and the method fails to arrive at a compact representation. This issue may be due to the relatively small penalties the method applies for constraining the optimal size of the mask and the entropy of the distribution. We plan to address this issue in the future by implementing more robust forms of regularization. In addition, although much of the GCN literature has focused on node features, more recent work also incorporates edge weights. In the context of our problem, edge weights could be assigned by using the Hi-C counts in the adjacency matrix. Another natural extension to our model would be to include other types of experimental data as features, such as promoter sequence or ATAC-seq measurements. Lastly, the GCN framework is flexible and general enough to be applied to many other classes of biological problems that require integrating diverse, multimodal data sets relationally.

In summary, GC-MERGE demonstrates proof-of-principle for using GCNs to predict gene expression using both local epigenetic features and long-range spatial interactions. More importantly, interpretation of this model allows us to propose plausible biological explanations of the key regulatory factors driving gene expression and provide guidance regarding promising hypotheses and new research directions.

## Acknowledgments

We are grateful to members of the COBRE-CBHD Computational Biology Core (CBC) at Brown University for helpful discussions and suggestions.

## Funding

Research reported in this publication was supported by an Institutional Development Award (IDeA) from the National Institute of General Medical Sciences of the National Institutes of Health under grant number P20GM109035.

## Disclosure

No competing financial interests exist.

## Supplementary Information

### S1#GC-MERGE model details

#### S1.1#Model architecture and training

The GC-MERGE architecture is represented in Figure 2. Here, the first layer of the model performs a graph convolution on the initial feature embeddings with an output embedding size of 256, followed by application of ReLU, a non-linear activation function. The second layer of the model performs another graph convolution with the same embedding size of 256 on the transformed representations, again followed by application of ReLU. Next, the output is fed into three successive linear layers of sizes 256, 256, and 2, respectively. A regularization step is performed by using a dropout layer with probability 0.5. The model was trained using ADAM, a stochastic gradient descent algorithm [Kingma and Ba, 2015]. We used the PyTorch Geometric package [Fey and Lenssen, 2019] to implement our code. Additional details regarding hyperparameter tuning can be found in the Supplemental Section S1.2.

#### S1.2#Hyperparameter tuning

Table S1 details the hyperparameters and the range of values we used to conduct a grid search to determine the optimized model. Specifically, we varied the number of graph convolutional layers, number of linear layers, embedding size for graph convolutional layers, linear layer sizes, and inclusion (or exclusion) of an activation function after the graph convolutional layers. Through earlier iterations of hyperparameter tuning, we also tested the type of activation functions used for the linear layers of the model (ReLU, LeakyReLU, sigmoid, or tanh), methods for accounting for background Hi-C counts, as well as dropout probabilities. Some combinations of hyperparameters were omitted from our grid search because the corresponding model’s memory requirements did not fit on the NVIDIA Titan RTX and Quadro RTX GPUs available to us on Brown University’s

**Table S1:** Hyperparameter combinations used for tuning in grid search. A grid search was conducted by varying the following hyperparameters: number of graph convolutional layers, number of linear layers, embedding size for graph convolutional layers, linear layer sizes, and inclusion/exclusion of activation function after the graph convolutional layers.

| Hyperparameter | Values |
| --- | --- |
| Number of graph convolutional layers | 1, 2 |
| Number of linear layers | 1, 2, 3 |
| Graph convolutional layer embedding sizes | 64, 128, 256, 384 |
| Linear layer sizes | Keep sizes of all linear layers constant;<br>alternatively, for each subsequent layer, divide size by 2 |
| Activation function after graph convolutional layers | Include; alternatively, do not include |

Center for Computation and Visualization (CCV) computing cluster. We recorded the loss curves for the training and validation sets over 800 epochs for the classification task and 1000 epochs for the regression task, by which time the model began to overfit. In addition, the data was split into sets of 70% for training, 15% for validation, and 15% for testing. The optimal hyperparameters for our final model that also proved to be computationally feasible are as follows: 2 graph convolutional layers, 3 linear layers, graph convolutional layer embedding size of 256, linear layer sizes that match that of the graph convolutional layers, and using an activation function (ReLU) after all graph convolutional layers and all linear layers except for the last.

**Figure S1:**
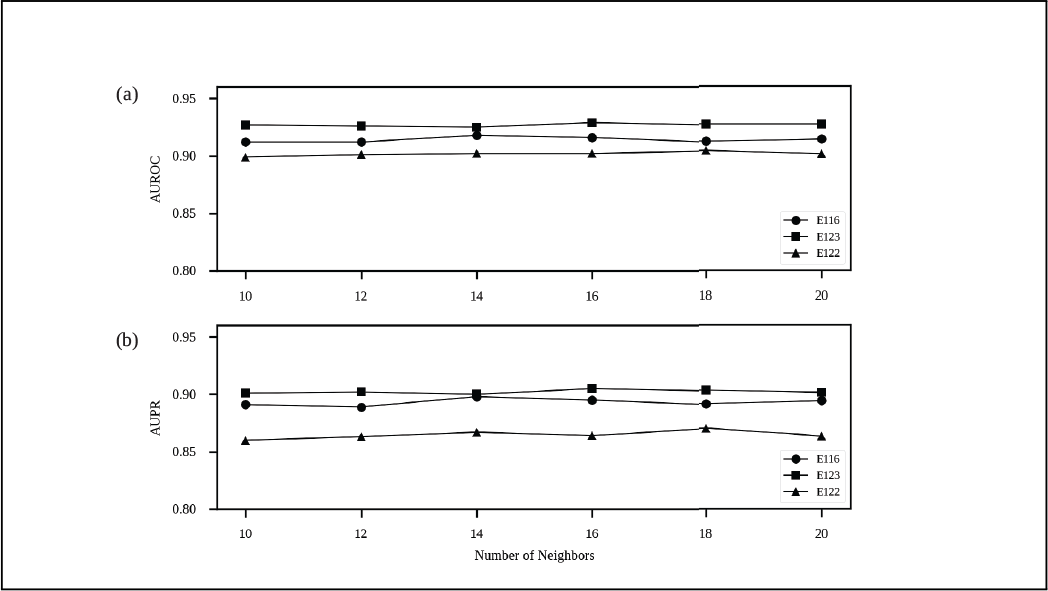
Effect of number of neighbors on classification performance. The performance of the model on the classification task is plotted as a function of the number of neighbors subsampled for each genic node. Including additional neighbors beyond 10 does not lead to substantial performance improvements with respect to either (a) AUROC or (b) AUPR.

**Figure S2:**
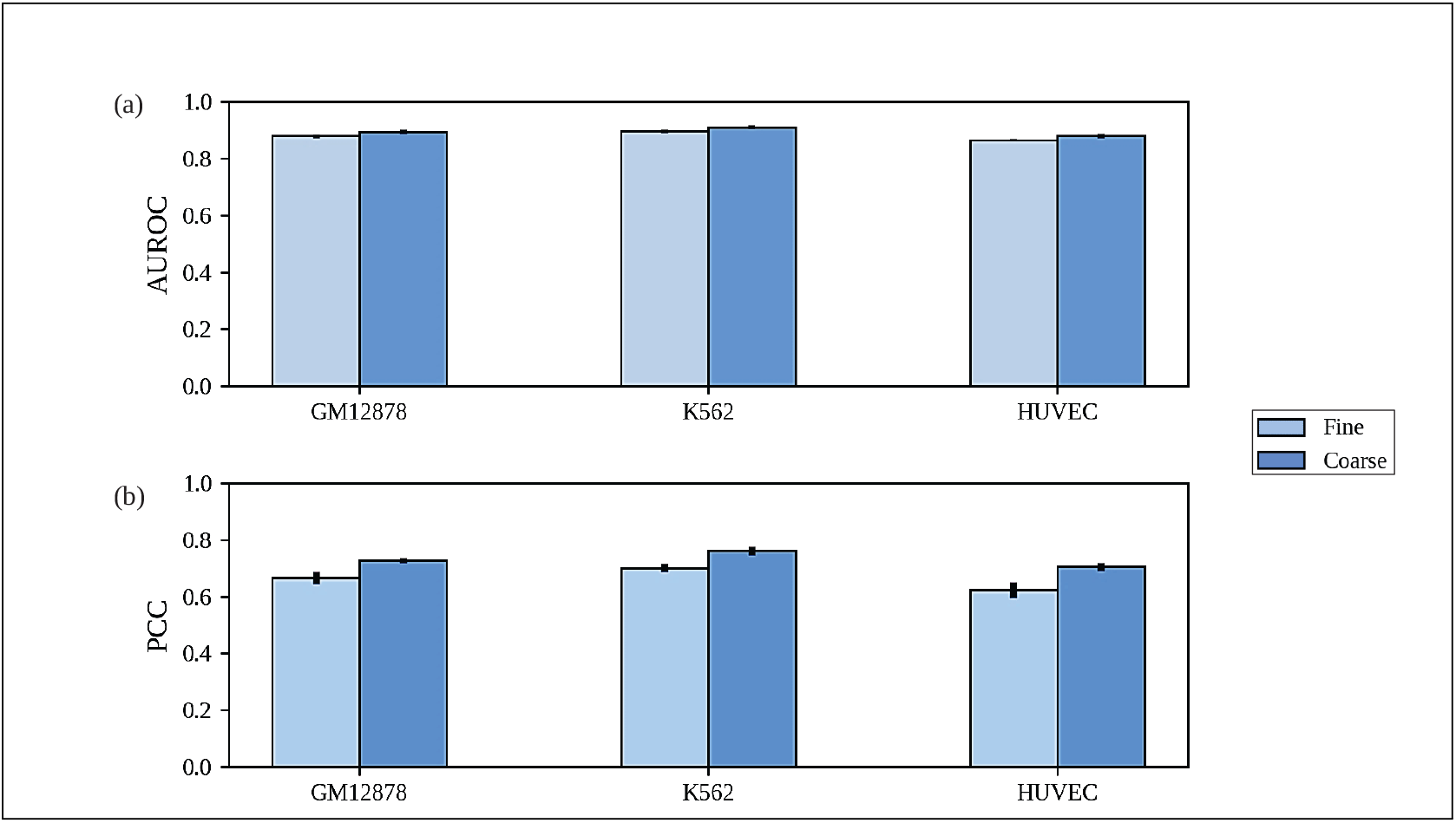
Comparison of fine-grained versus coarse-grained ChIP-seq signals. For the coarse-grained resolution, ChIP-seq signals were averaged over the entire Hi-C bin (10000 bp resolution). For the fine-grained resolution, ChIP-seq signals were first averaged over 1000 bp bins and then fed into two embedding linear layers followed by ReLU. The output of these embedding layers was then was used to feature annotate each node. (a) For the classification task, the fine-grained resolution ChIP-seq data performs slightly worse than or comparable to that of the coarse-grained resolution ChIP-seq data as measured by AUROC. (b) For the regression task, the fine-grained resolution ChIP-seq data produces performance worse than or comparable to the coarse-grained resolution ChIP-seq data as measured by PCC.

### S2#Analysis of exemplar genes

• **SIDT1** encodes a transmembrane dsRNA-gated channel protein and is part of a larger family of proteins necessary for systemic RNA interference [Elhassan et al., 2012, Entrez Gene, 1988]. This gene has also been implicated in chemoresistance to the drug gemcitabine in adenocarcinoma cells [Elhassan et al., 2012].
• **AKR1B1** encodes an enzyme that belongs to the aldo-keto reductase family. It has also been identified as a key player in complications associated with diabetes [Donaghue et al., 2005, Entrez Gene, 1988].
• **LAPTM5** encodes a receptor protein that spans the lysosomal membrane [Entrez Gene, 1988]. It is highly expressed in immune cells and plays a role in the downregulation of T and

**Table S2:** Node coordinates for all exemplar genes: SIDT1, AKR1B1, LAPTM5, TOP2B, BAX, HNRNPA1, PRDX2, and RAD23A. For each gene, the second and third columns list the corresponding node identifiers and the chromosome coordinates, respectively. The fourth column lists the gene’s actual chromosomal coordinates. Note that the transcription start site was used as the basis for assigning each gene to a node.

| Gene | Node ID | Coordinates |
| --- | --- | --- |
| TOP2B | 51806 | chr3:25639475-25706398 |
| SIDT1 | 60561 | chr3:113251143-113348425 |
| LAPTM5 | 3123 | chr1:31205316-31230667 |
| AKR1B1 | 136736 | chr7:134127127-134144036 |
| BAX | 264956 | chr19:49458072-49465055 |
| HNRNPA1 | 200470 | chr12:54673977-54680872 |
| PRDX2 | 261302 | chr19:12907634-12912694 |
| RAD23A | 261316 | chr19:13056654-13064448 |

B cell receptors and the upregulation of macrophage cytokine production [Glowacka et al., 2012].

• **TOP2B** encodes DNA topoisomerase II beta, a protein that controls the topological state of DNA during transcription and replication [Entrez Gene, 1988]. It transiently breaks and then reforms duplex DNA, relieving torsional stress. Mutations in this enzyme can lead to B cell immunodeficiency [Broderick et al., 2019].
• **BAX** encodes a protein that forms a heterodimeric complex with BCL2, which activates apoptosis by aggregating at the mitochondrial membrane and inducing its permeabilization [Entrez Gene, 1988, Peña-Blanco and Garćıa-Sáez, 2018]. The tumor suppressor gene P53 plays a role in its regulation.
• **HNRNPA1** encodes a protein that forms part of the heterogeneous nuclear ribonucleoprotein (hnRNP) complex, which binds to nuclear pre-mRNA and helps to regulate RNA transport, metabolism, and splicing [Entrez Gene, 1988, Roy et al., 2017]. Mutations in this gene have been linked to the development of amyotrophic lateral sclerosis.
• **PRDX2** encodes an enzyme that reduces hydrogen peroxide and alkyl hydroperoxides [Entrez Gene, 1988]. It protects against oxidative stress [Jin et al., 2020] as well as stabilizes hemoglobin, making it a therapeutic target for the treatment of hemolytic anemia.
• **RAD23A** encodes a protein that carries out nucleotide excision repair [Fang et al., 2013, Entrez Gene, 1988]. It also plays a role in transporting poly-ubiquitinated proteins to the proteasome for degradation.

**Table S3:**
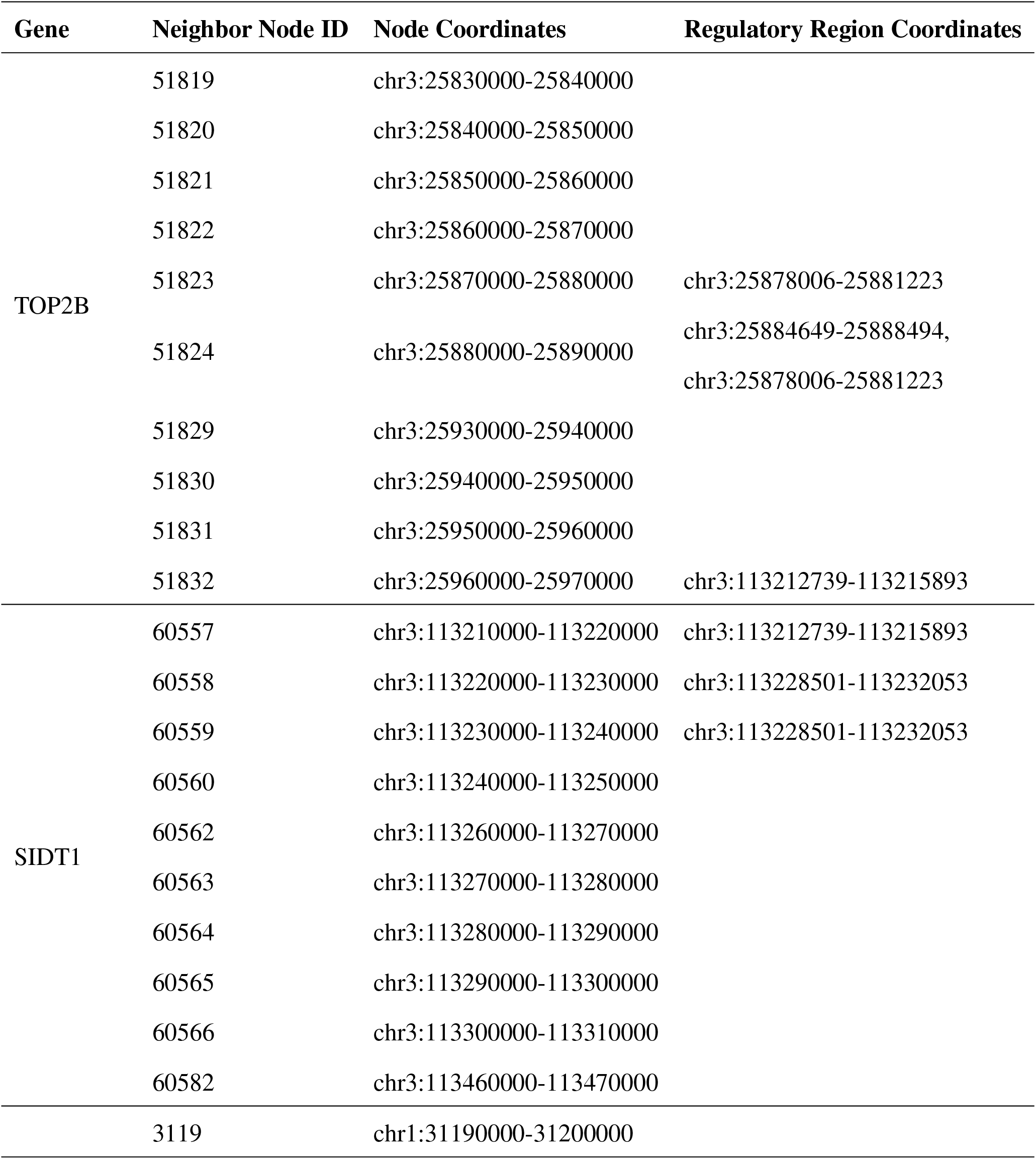
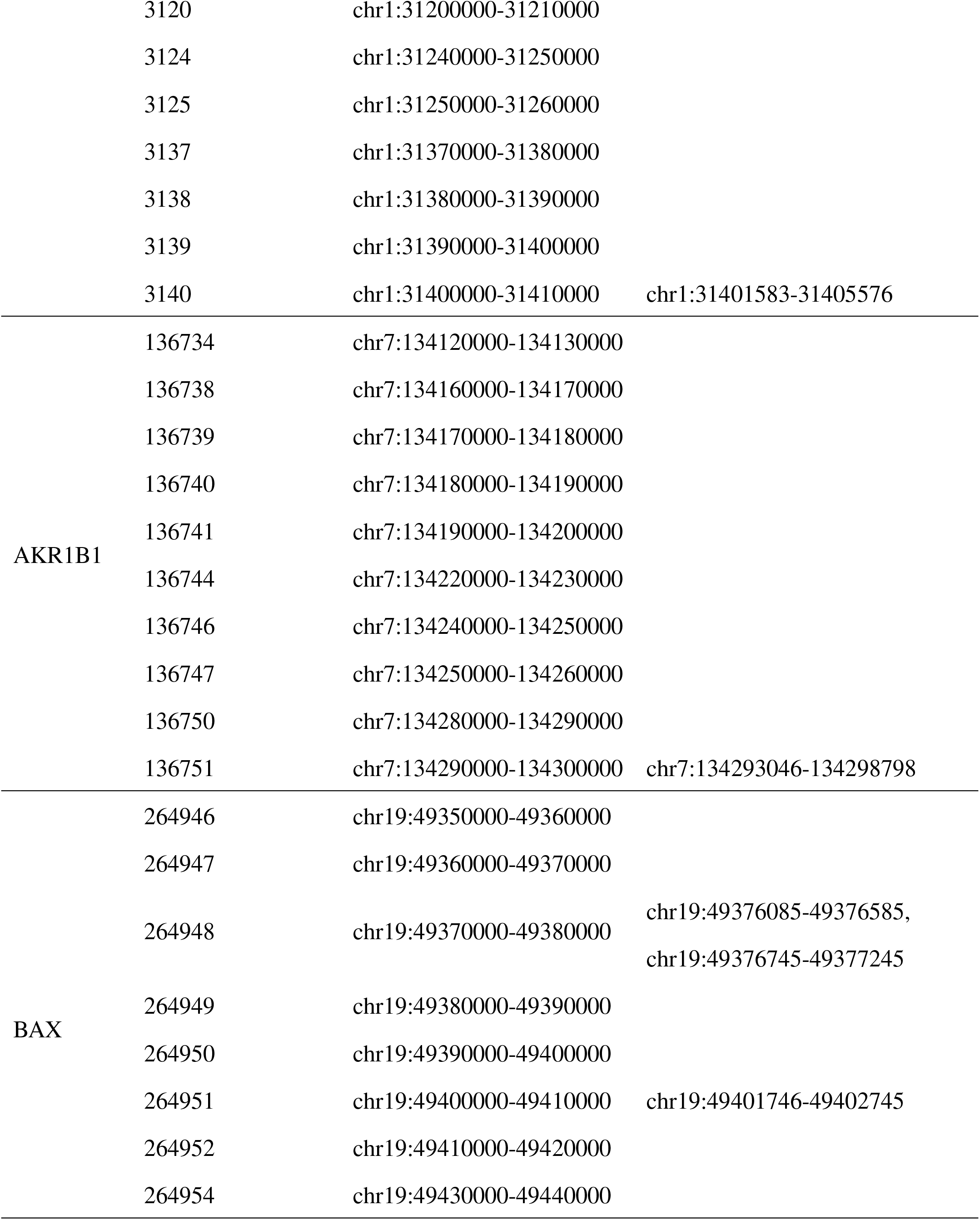
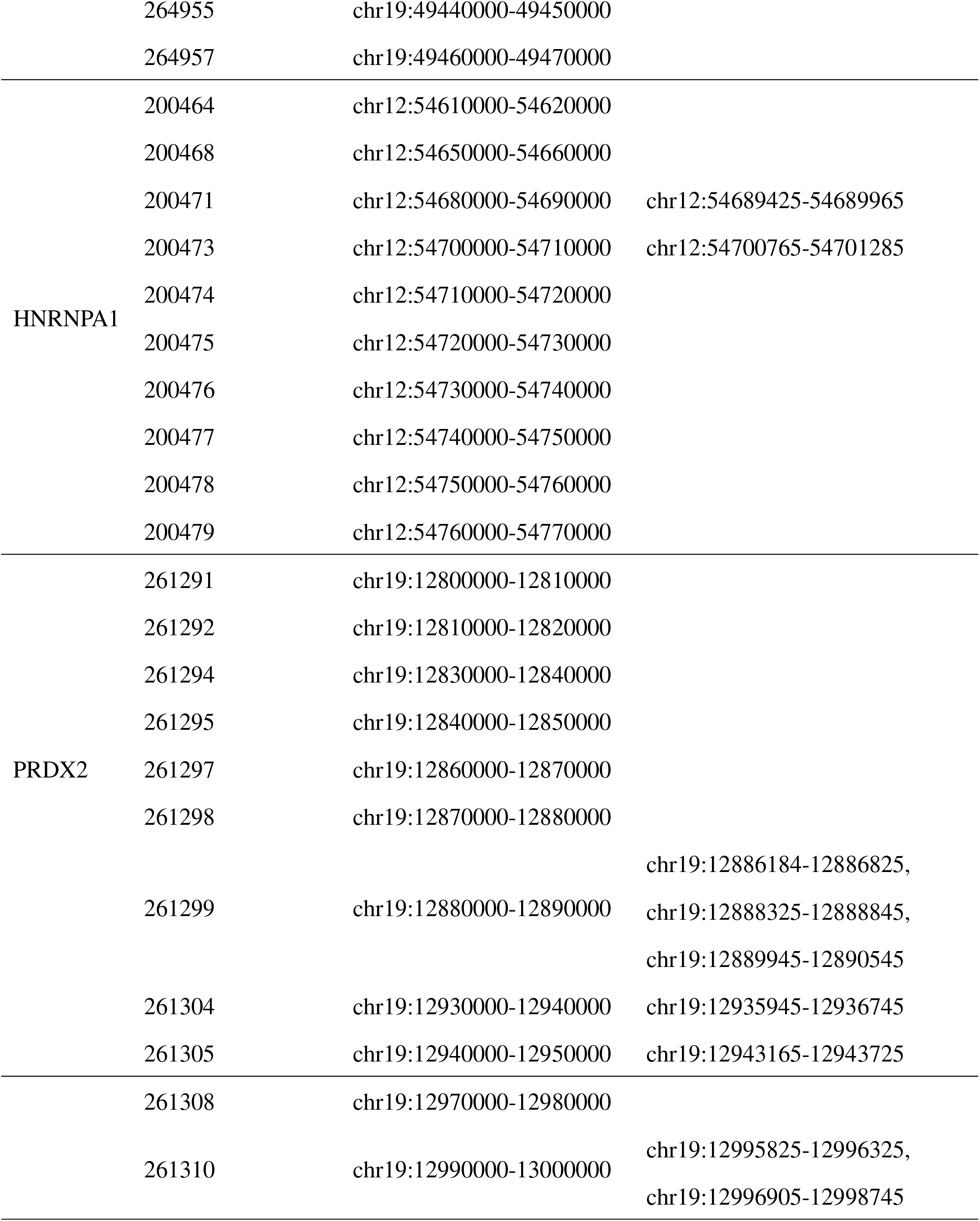
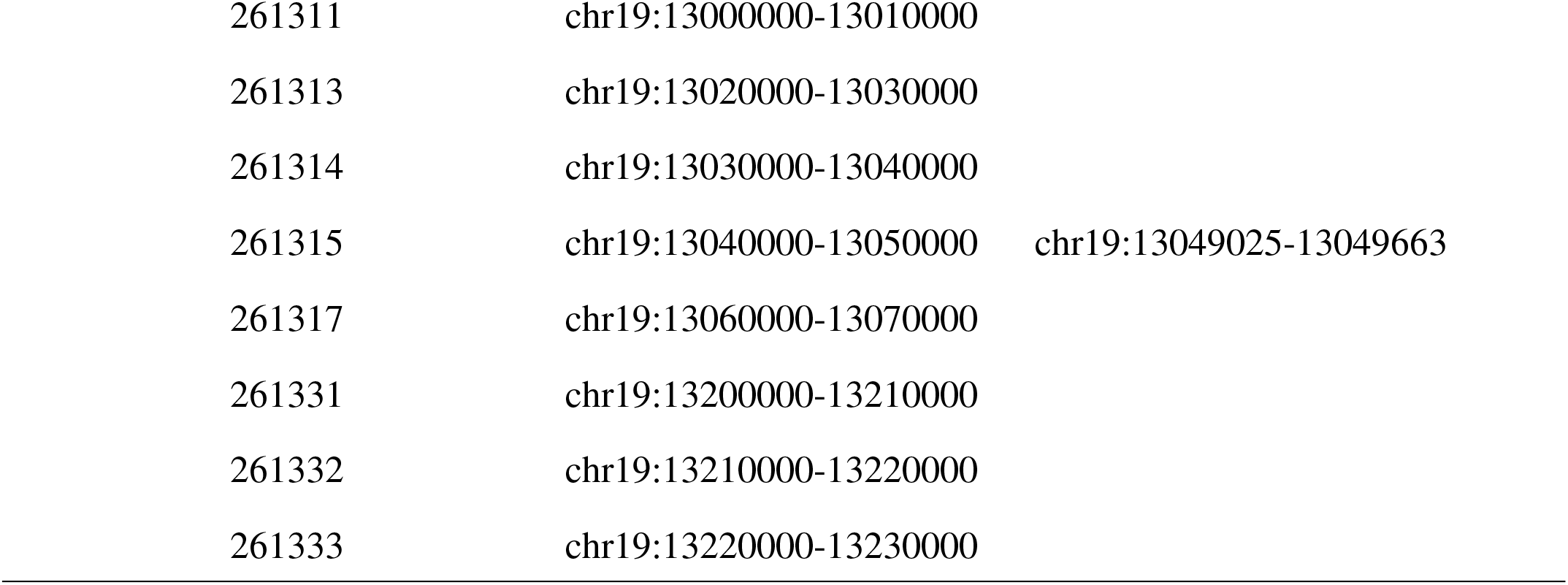
Neighbor coordinates for exemplar genes. The second column lists the node identifiers for all neighboring nodes of the relevant gene, including neighboring nodes that contain interacting fragments as well as those that do not. The third column third lists the corresponding chromosome coordinates for the node identifier. The fourth column lists the regulatory fragments that interact with each gene as described in Jung et al. [2019] and Fulco et al. [2019]

**Figure S3:**
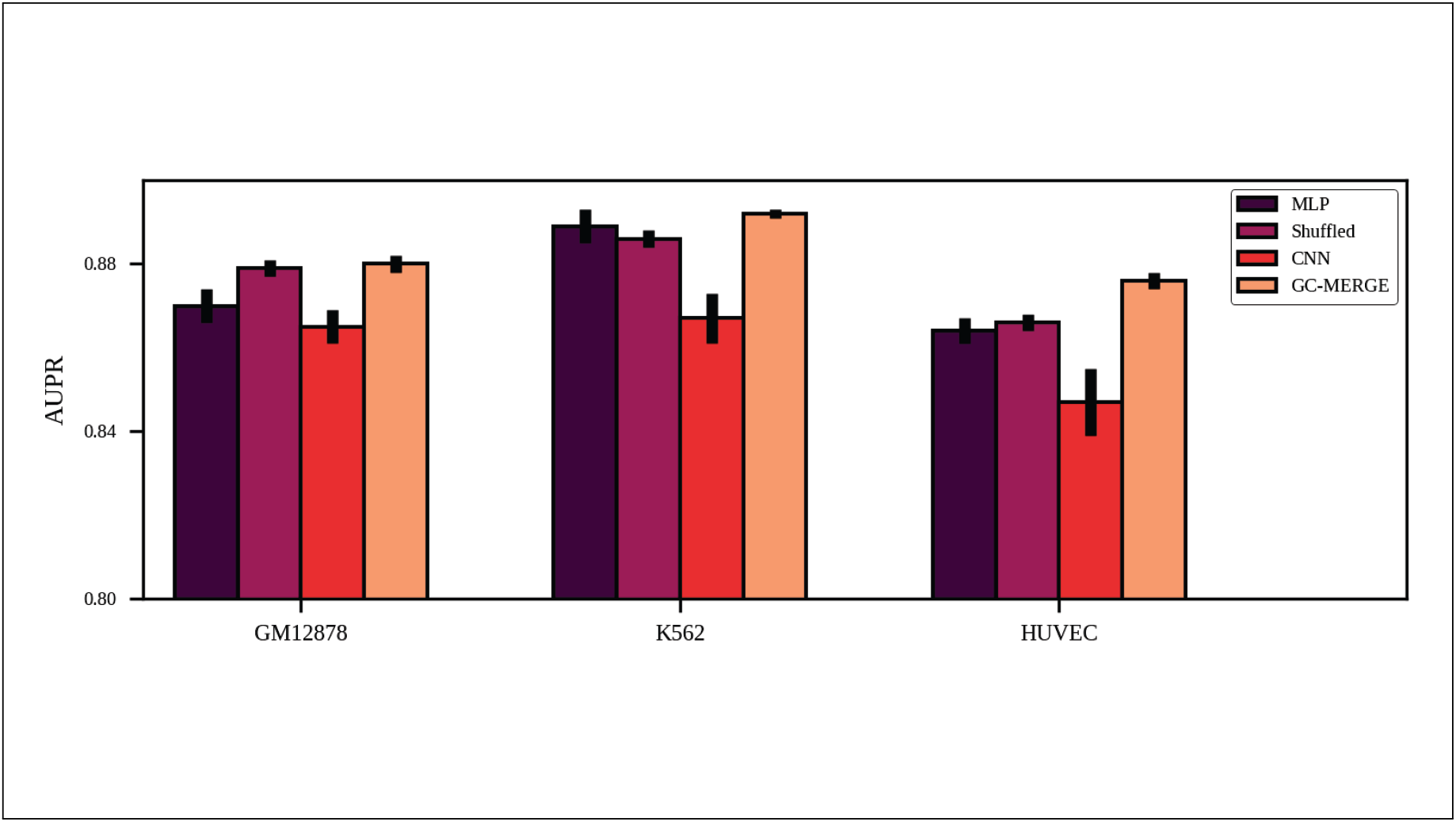
Comparison of AUPR scores for all models. GC-MERGE gives state-of-the-art performance for classifying genes as either having high expression or low expression. Using the AUPR metric, GC-MERGE obtains scores of 0.88, 0.89, and 0.88 for GM12878, K562, and HUVEC, respectively.

**Figure S4:**
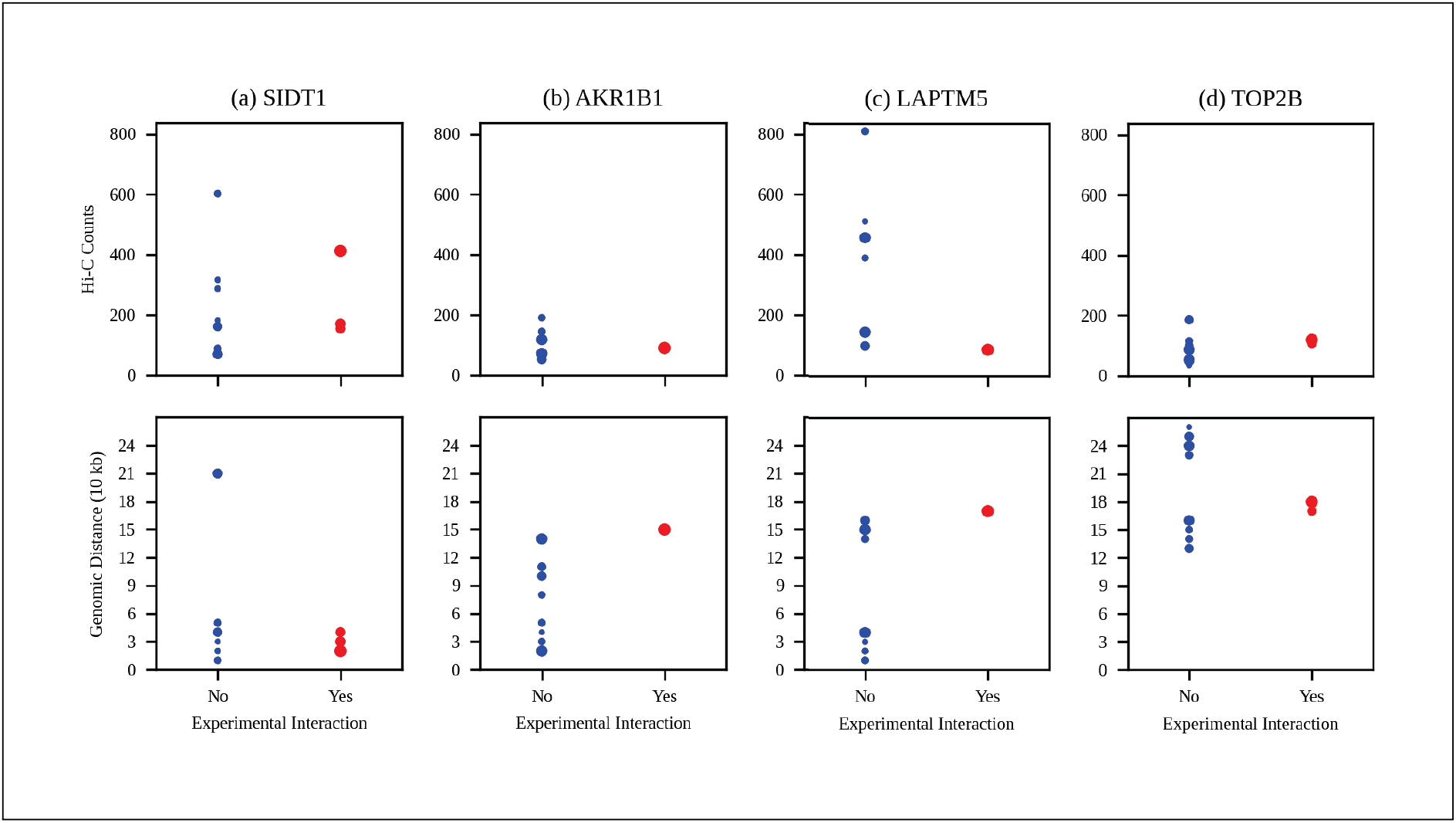
Relationships among importance scores, genomic distances, and Hi-C counts for all exemplar genes with promoter-capture Hi-C validated regulatory interactions. The exemplar genes are shown by column as follows: (a) TOP2B, (b) SIDT1, (c) LAPTM5, and (d) AKR1B1. The size of each data point corresponds to the neighbor node’s scaled importance score. Nodes corresponding to experimentally validated interacting fragments are denoted in red and all others are denoted in blue. The top panel plots Hi-C counts classified according to experimental validation, while as the bottom panel plots genomic distance versus experimental interaction. Neither Hi-C counts nor genomic distance correlate with experimentally validated interactions.

**Table S4:** Relationships among node importance scores, distance from target gene, and Hi-C counts. For each exemplar gene, the the node IDs of its neighbors are shown (second column) along with their importance scores (third column), distances from the target gene node (fourth column), and normalized Hi-C frequency counts (fifth column).

| Gene | Neighbor Node ID | Importance Score | Distance (10 kb) | Hi-C Counts |
| --- | --- | --- | --- | --- |
| TOP2B | 51819 | 4.3 | 13 | 186 |
|  | 51820 | 3.0 | 14 | 116 |
|  | 51821 | 2.5 | 15 | 97 |
|  | 51822 | 6.9 | 16 | 87 |
|  | 51823 | 4.5 | 17 | 105 |
|  | 51824 | 10.0 | 18 | 120 |
|  | 51829 | 3.5 | 23 | 101 |
|  | 51830 | 6.8 | 24 | 54 |
|  | 51831 | 5.3 | 25 | 47 |
|  | 51832 | 1.0 | 26 | 34 |
| SIDT1 | 60557 | 5.7 | 4 | 155 |
|  | 60558 | 6.6 | 3 | 171 |
|  | 60559 | 10.0 | 2 | 413 |
|  | 60560 | 1.3 | 1 | 316 |
|  | 60562 | 2.7 | 1 | 603 |
|  | 60563 | 1.6 | 2 | 287 |
|  | 60564 | 1.0 | 3 | 182 |
|  | 60565 | 4.7 | 4 | 162 |
|  | 60566 | 2.9 | 5 | 88 |
|  | 60582 | 5.6 | 21 | 70 |
| LAPTM5 | 3119 | 7.9 | 4 | 459 |
|  | 3120 | 1.0 | 3 | 513 |
|  | 3124 | 3.0 | 1 | 812 |
|  | 3125 | 2.0 | 2 | 391 |
|  | 3137 | 3.2 | 14 | 101 |
|  | 3138 | 7.8 | 15 | 145 |
|  | 3139 | 5.1 | 16 | 99 |
|  | 3140 | 10.0 | 17 | 86 |
| AKR1B1 | 136734 | 2.5 | 2 | 191 |
|  | 136738 | 7.7 | 2 | 118 |
|  | 136739 | 2.5 | 3 | 65 |
|  | 136740 | 1.0 | 4 | 78 |
|  | 136741 | 2.8 | 5 | 145 |
|  | 136744 | 2.4 | 8 | 63 |
|  | 136746 | 5.0 | 10 | 53 |
|  | 136747 | 4.0 | 11 | 66 |
|  | 136750 | 7.7 | 14 | 73 |
|  | 136751 | 10.0 | 15 | 90 |
| BAX | 264946 | 3.9 | 10 | 38 |
|  | 264947 | 4.6 | 9 | 45 |
|  | 264948 | 8.4 | 8 | 61 |
|  | 264949 | 3.5 | 7 | 69 |
|  | 264950 | 3.0 | 6 | 55 |
|  | 264951 | 10.0 | 5 | 79 |
|  | 264952 | 1.0 | 4 | 103 |
|  | 264954 | 3.3 | 2 | 152 |
|  | 264955 | 7.4 | 1 | 265 |
|  | 264957 | 6.7 | 1 | 153 |
| HNRNPA1 | 200464 | 4.8 | 6 | 44 |
|  | 200468 | 1.6 | 2 | 67 |
|  | 200471 | 6.6 | 1 | 83 |
|  | 200473 | 10.0 | 3 | 27 |
|  | 200474 | 1.1 | 4 | 79 |
|  | 200475 | 5.8 | 5 | 77 |
|  | 200476 | 1.0 | 6 | 48 |
|  | 200477 | 3.3 | 7 | 37 |
|  | 200478 | 3.9 | 8 | 79 |
|  | 200479 | 7.6 | 9 | 49 |
| PRDX2 | 261291 | 1.0 | 11 | 34 |
|  | 261292 | 3.3 | 10 | 44 |
|  | 261294 | 4.8 | 8 | 57 |
|  | 261295 | 3.6 | 7 | 47 |
|  | 261297 | 4.5 | 5 | 36 |
|  | 261298 | 5.2 | 4 | 70 |
|  | 261299 | 10.0 | 3 | 79 |
|  | 261304 | 8.1 | 2 | 116 |
|  | 261305 | 5.7 | 3 | 117 |
| RAD23A | 261308 | 5.5 | 8 | 47 |
|  | 261310 | 9.8 | 6 | 90 |
|  | 261311 | 7.6 | 5 | 62 |
|  | 261313 | 4.9 | 3 | 32 |
|  | 261314 | 1.0 | 2 | 73 |
|  | 261315 | 10.0 | 1 | 162 |
|  | 261317 | 4.3 | 1 | 122 |
|  | 261331 | 2.8 | 15 | 32 |
|  | 261332 | 5.7 | 16 | 20 |
|  | 261333 | 8.1 | 17 | 26 |

**Figure S5:**
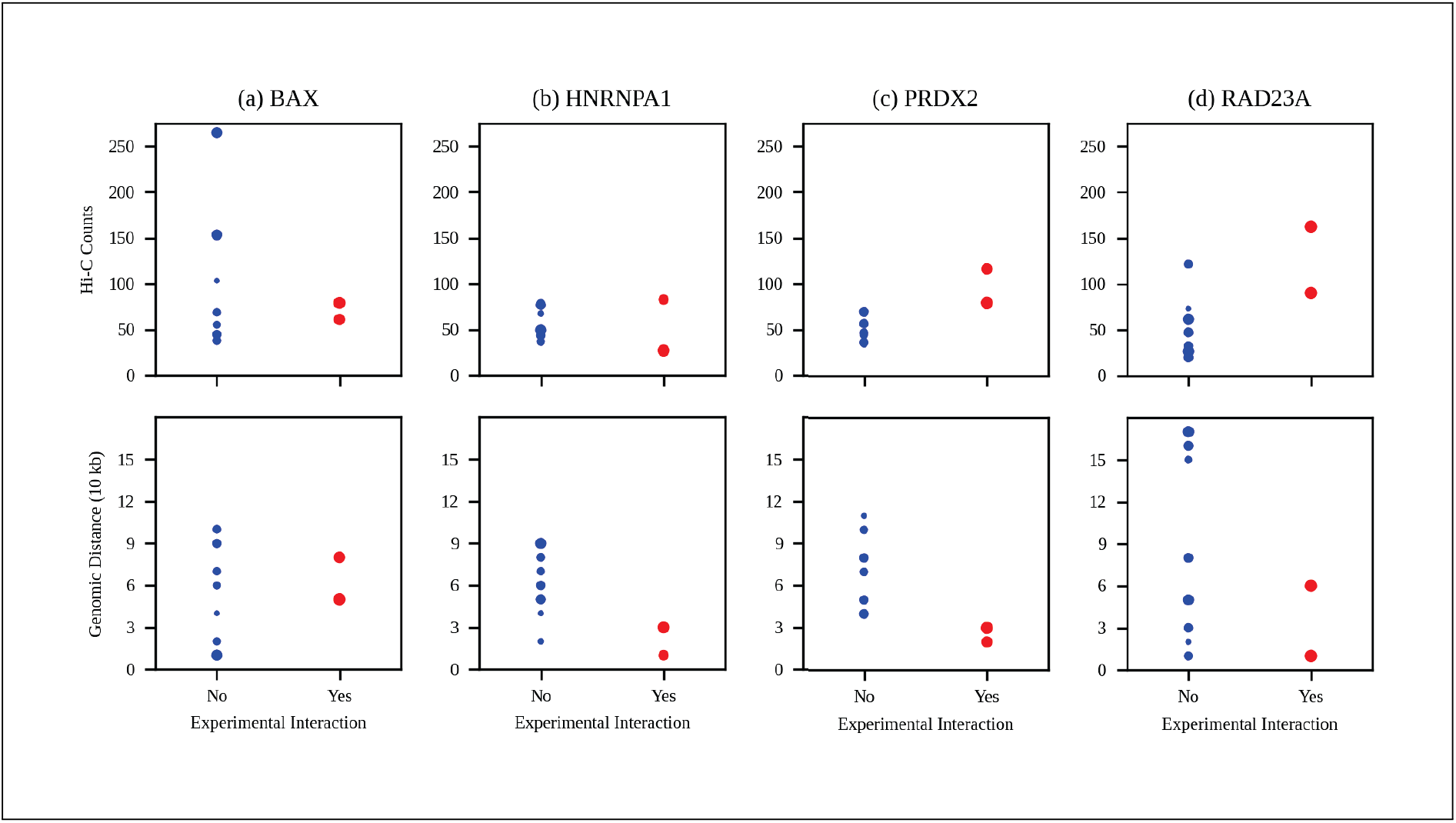
Relationships among importance scores, genomic distances, and Hi-C counts for all exemplar genes with CRISPRi-FlowFISH validated regulatory interactions. The exemplar genes are shown by column as follows: (a) BAX, (b) HNRNPA1, (c) PRDX2, and (d) RAD23A. The size of each data point corresponds to the neighbor node’s scaled importance score. Nodes corresponding to experimentally validated interacting fragments are denoted in red and all others are denoted in blue. The top panel plots Hi-C counts classified according to experimental validation, while as the bottom panel plots genomic distance versus experimental interaction. Neither Hi-C counts nor genomic distance correlate with experimentally validated interactions.

## Notes

### Competing Interest Statement

The authors have declared no competing interest.

### Summary of Updates

Added new results, revised descriptions

